# Modulation of human intracranial theta oscillations during freely moving spatial navigation and memory

**DOI:** 10.1101/738807

**Authors:** Zahra M. Aghajan, Diane Villaroman, Sonja Hiller, Tyler J. Wishard, Uros Topalovic, Leonardo Christov-Moore, Nader Shaterian, Nicholas R. Hasulak, Barbara Knowlton, Dawn Eliashiv, Vikram Rao, Itzhak Fried, Nanthia Suthana

**Affiliations:** Department of Psychiatry and Biobehavioral Sciences, Jane and Terry Semel Institute for Neuroscience and Human Behavior, University of California Los Angeles, Los Angeles, CA; Fab Lab Connect, Golden Beach, FL; NeuroPace Inc., Mountain View, CA; Department of Psychology, University of California Los Angeles, Los Angeles, CA; Department of Neurology, University of California Los Angeles, Los Angeles, CA; Department of Neurology, University of California San Francisco, San Francisco, CA; Department of Neurosurgery, David Geffen School of Medicine, University of California Los Angeles, Los Angeles, CA; Functional Neurosurgery Unit, Tel Aviv Medical Center and Sackler School of Medicine, Tel Aviv University, Tel Aviv, Israel

## Abstract

How the human brain supports accurate navigation of a learned environment has been an active topic of research for nearly a century^1–5^. In rodents, the theta rhythm within the medial temporal lobe (MTL) has been proposed as a neural basis for fragmenting incoming information and temporally organizing experiences and is thus widely implicated in spatial and episodic memory^6^. In addition, high-frequency theta (~8Hz) is associated with navigation, and loss of theta results in spatial memory deficits in rats ^7^. Recently, high-frequency theta oscillations during ambulatory movement have been identified in humans^8,9^, though their relationship to spatial memory remains unexplored. Here, we were able to record MTL activity during spatial memory and navigation in freely moving humans immersed in a room-scale virtual reality (VR) environment. Naturalistic movements were captured using motion tracking combined with wireless VR in participants implanted with an intracranial electroencephalographic (iEEG) recording system for the treatment of epilepsy. We found that prevalence of theta oscillations across brain sites during both learning and recall of spatial locations during ambulatory navigation is critically linked to memory performance. This finding supports the reinstatement hypothesis of episodic memory—thought to underlie our ability to recreate a prior experience^10–12^—and suggests that theta prevalence within the MTL may act as a potential representational state for memory reinstatement during spatial navigation. Additionally, we found that theta power is hexadirectionally modulated^13–15^ as a function of the direction of physical movement, most prominently after learning has occurred. This effect bears a resemblance to the rodent grid cell system^16^ and suggests an analog in human navigation. Taken together, our results provide the first characterization of neural oscillations in the human MTL during ambulatory spatial memory tasks and provide a platform for future investigations of neural mechanisms underlying freely moving navigation in humans.

## Main

Five participants (one congenitally blind, Extended Data Table 1) chronically implanted with the NeuroPace RNS^®^ System (for treatment of refractory focal epilepsy) performed ambulatory virtual spatial navigation (VR) tasks (Fig. 1a, Extended Data Fig. 1). The RNS System was used to record iEEG from depth electrode contacts^8^ (Fig. 1a, Extended Data Fig. 2). VR tasks were implemented in the Unity game engine and were configured to communicate with a Motion Capture system via WiFi. The physical movement of the participant in the real room was mapped to body position in the virtual space such that the scene was updated according to their motion in a one-to-one manner (Methods). Participants were instructed to learn the location and order of multiple virtual translucent yellow cylinders (henceforth referred to as halos; Fig. 1a) during a learning (encoding) phase. After a non-mnemonic distractor task, the recall (retrieval) phase began during which participants navigated to previously learned locations in their original order, pressing a joystick button to indicate when they arrived within each halo. There were two variations in how the halos were presented during encoding: 1) Directed navigation: halos appeared one at a time and remained visible until the participant reached them (Supplementary Video 1); 2) Discovery navigation: halos remained hidden and participants had to freely maneuver through the room until they reached a halo, at which point it appeared (Supplementary Video 2). Order of the memory tasks (directed vs. discovery) was counterbalanced across the visually sighted participants. For the blind participant, auditory versions of the spatial memory tasks were completed. Specifically, the proximity of the participant to the halos was reflected by the frequency of the sound (i.e., higher frequency and faster beeping indicated getting closer). The discovery navigation task was completed first in the blind participant. For all participants, the directed and discovery navigation memory task had 10 and 6 predetermined locations respectively, in order to match the difficulty levels of the task; the number of locations were determined based on prior pilot testing in normal healthy participants (Fig. 1a-b; Methods).

**Figure 1.**
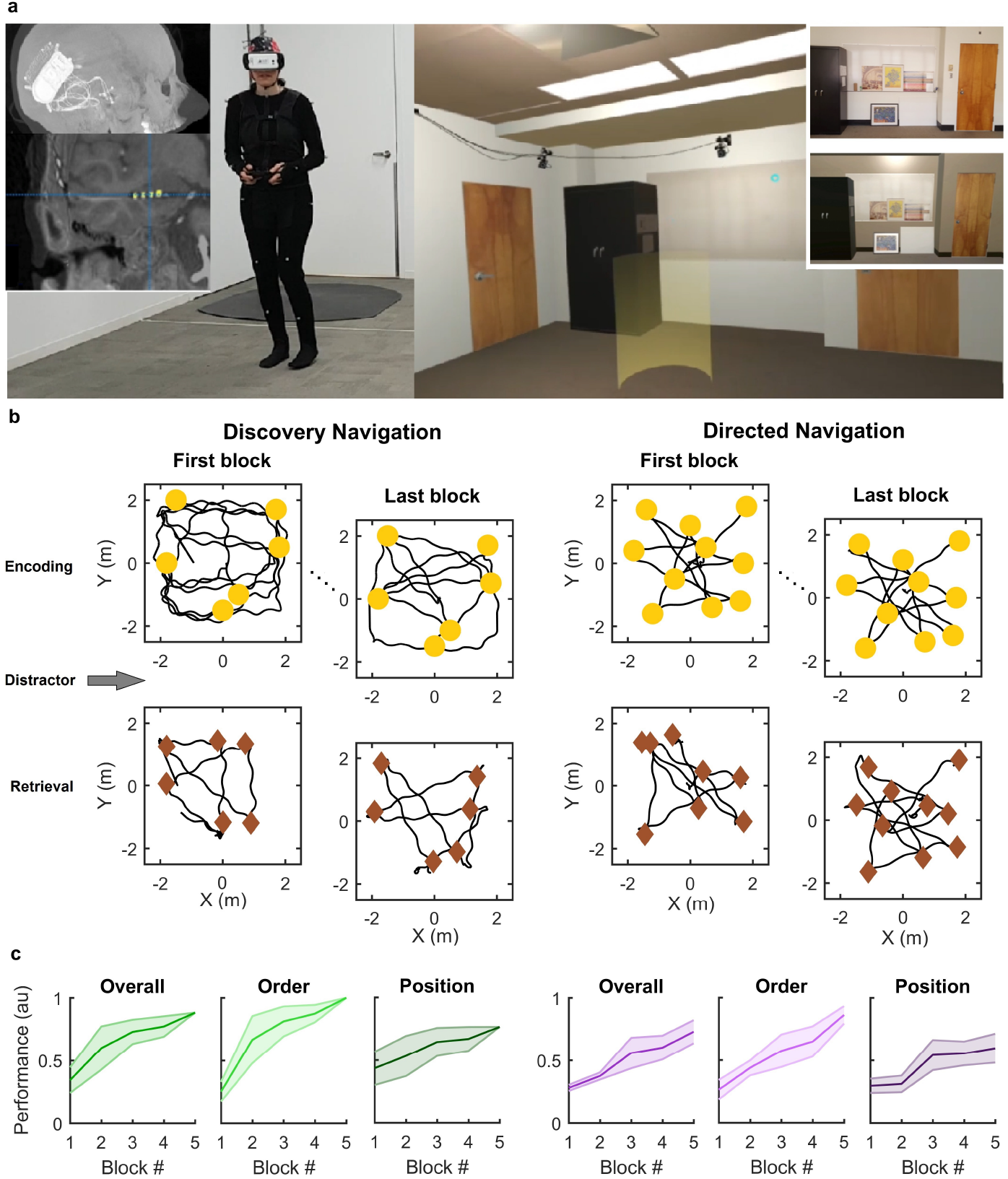
Schematic of ambulatory virtual spatial navigation tasks and participants’ behavioral performance. **a)** Left - Shown is an example CT (lateral view) and MRI (coronal view) from one participant implanted with the wireless NeuroPace RNS System and the wireless VR headset with joystick setup. Right - Example view of a participant’s view through the VR headset (inset: a photograph of separate never before seen room of the same size dimensions [top] from which a virtual rendering was created [bottom]). **b)** Example behavior of participant 1 (P1) during discovery navigation (left) and directed navigation (right) during the first and last of 5 blocks. Both the directed and discovery navigation tasks consisted of encoding, distraction, and retrieval phases within each block. During encoding, participants walked through the room (trajectory shown in black) and were asked to remember the location and order of multiple yellow virtual translucent cylinders (halos, yellow circles; also visible in **(a)**). During recall, participants navigated to the locations they remembered and indicated their arrival by pressing a joystick button (brown diamonds). **c)** Behavioral performance of all sighted participants (n=4; mean ± s.e.m) during discovery (left; green shades) and directed (right, purple) navigation tasks is shown across blocks. The overall performance measure includes both order and position components.

Behavioral performance was quantified in terms of position (the proximity of the recalled location to the original halo location) and order (the recalled order of halos compared to the original order) and combined into a composite score (Fig. 1b, Methods). Participants were able to successfully learn and recall spatial locations with on average, performance improving throughout the entire session of multiple blocks (Fig. 1c). The congenitally blind participant was excluded from the presented behavioral analysis.

Next, we asked whether movement-related theta oscillations^8,9,17^ could be observed in the iEEG data (Fig 2a) during the two navigation tasks. Indeed, high frequency (6-12Hz) theta was present in all participants as can be seen both in the raw traces (Fig. 2b) as well as in the spectral power analysis (Fig. 2c; Methods). We hypothesized that theta may exhibit different properties during discovery versus directed navigation as the former presumably involves a more exploratory approach during encoding and may, thus, recruit more theta oscillations^18,19^. Thus, we quantified the prevalence of theta (percentage of time with significant oscillations) in each task—using previous methods^8,20,21^—during encoding (Fig. 3d) and retrieval (Extended Data Fig. 3) independently. To our surprise, we did not find significant differences in theta prevalence between discovery and directed navigation in any of the participants except for one participant (Fig. 2d; P3: *p* < 0.05, cluster-based permutation test; other participants: *p*>0.05). These findings suggest that other factors, beyond the degree of exploratory behavior, may be influencing how often theta oscillations are present.

**Figure 2.**
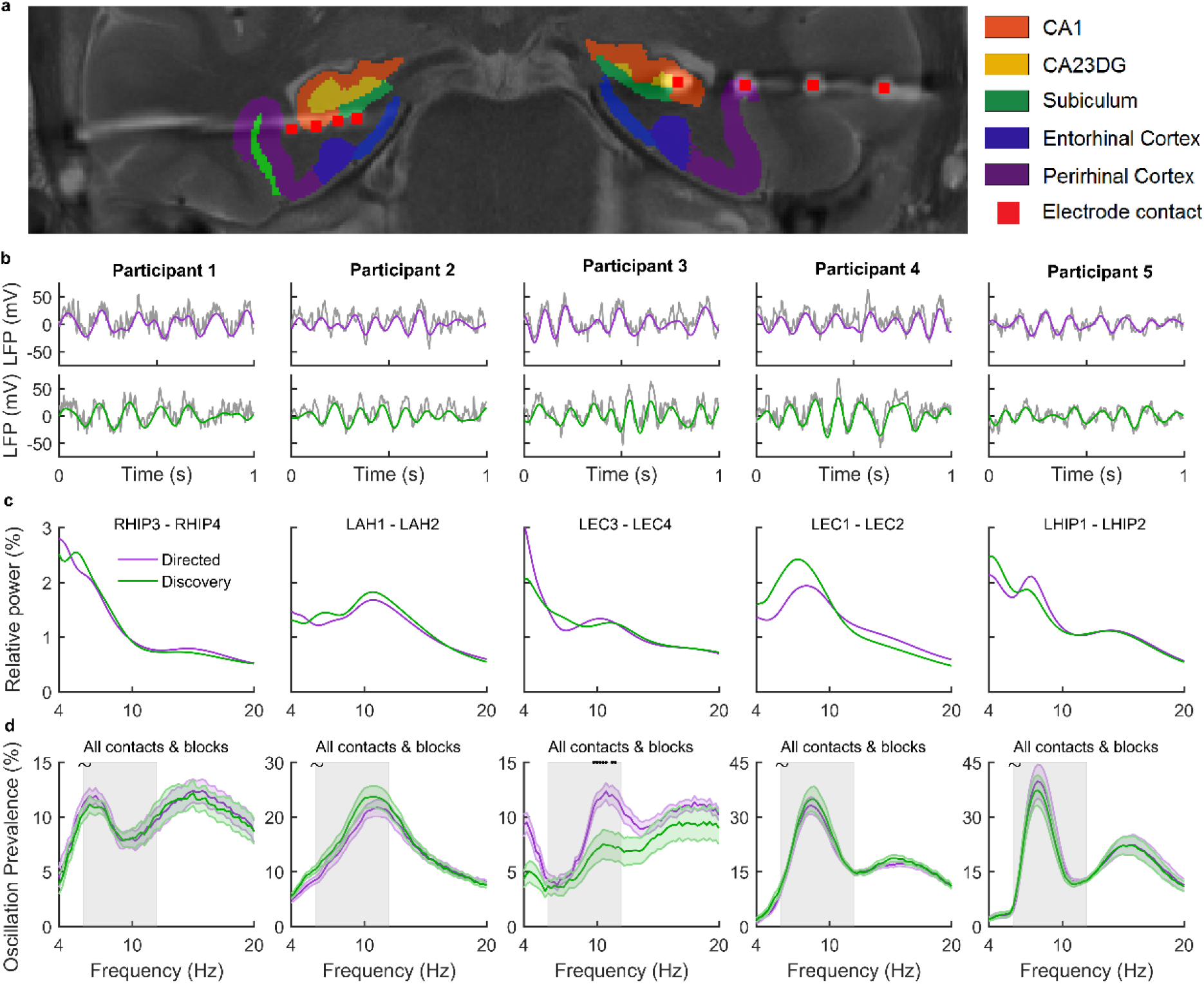
Presence and similar prevalence of theta oscillations during directed and discovery spatial navigation memory tasks. **a)** Sample electrode locations from an example participant (P4) overlaid onto the coronal pre-operative high-resolution MRI. MTL subregions created from automated segmentation are shown, with different colors corresponding to different subregions (Methods). **b)** Example one-second-long raw iEEG traces (gray) from representative recording channels overlaid with filtered (6-12Hz) theta oscillations (directed and discovery navigation shown in purple and green respectively). Each column corresponds to a single participant. The exact location of the recording contacts shown here are as follows (Extended Data Table 2): P1: CA1; P2: CA23DG-CA23DG; P3: CA23DG-CA23DG/CA1; P4: Hippocampus-Perirhinal Cortex; P5: CA1-CA1. **c)** Example (normalized) power spectra from representative channels in each participant demonstrating the presence of theta oscillations (peaks in the 6-12Hz) during directed (purple) and discovery (green) navigation. **d)** Theta prevalence was similar (~*p*>0.05; clustered-based permutation test) between directed and discovery navigation in all participants with the exception of participant 3 (*p*<0.05; clustered-based permutation test; black dots correspond to frequencies with significant differences in theta prevalence). In each participant, this was done when considering all channels and behavioral blocks (n1 and n2 correspond to the number of data points during directed and discovery navigation respectively; P1: n1=20, n2=16; P2: n1=24, n2=20; P3: n1=20, n2=16; P1: n1=20, n2=16; P1: n1=20, n2=16). Shown are the mean (dark colored lines) ± s.e.m values (shaded color areas). The gray shaded area corresponds to the theta frequency band (6-12Hz) of interest.

**Figure 3.**
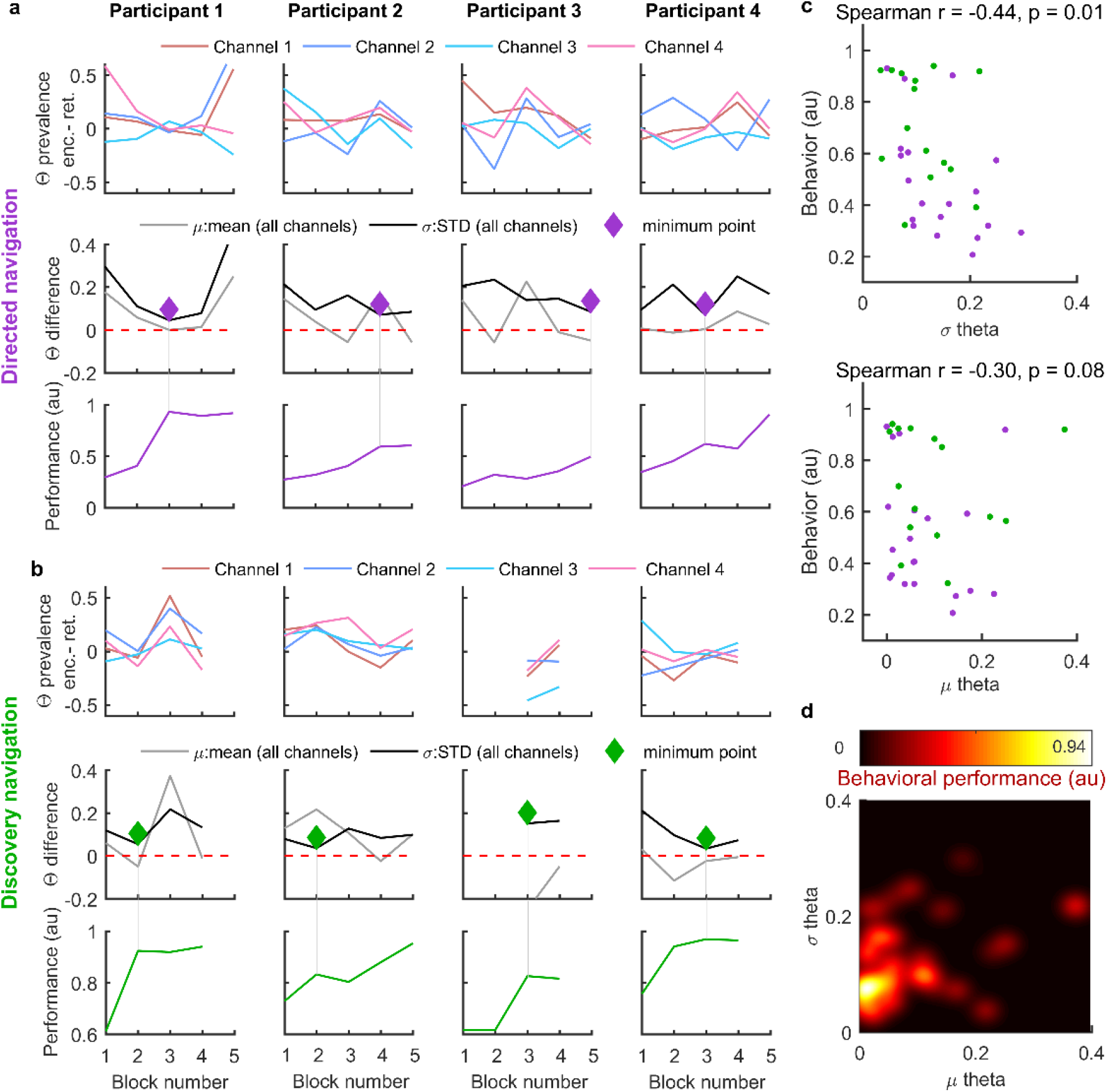
Similar patterns of theta prevalence during encoding and retrieval coincided with increased memory performance. **a)** Directed navigation; Top: Difference in theta prevalence between encoding and retrieval (normalized by the sum) for all recording contacts (different colors) across behavioral blocks for each participant (different columns). Middle: The mean (gray; μ) and standard deviation (black; σ) of theta difference (henceforth referred to as μ theta and σ theta respectively) across channels (shown above). Diamonds correspond to the block within each participant where the minimum standard deviation of theta difference across channels was detected. Bottom: Behavioral performance across recall blocks. Vertical gray lines indicate the correspondence between behavioral blocks and minimum theta difference. **b)** Same as in (a) but for discovery navigation. *iEEG data from participant 3 was only captured during retrieval phases of blocks 3-4. **c)** Top: Behavioral performance within each block was negatively correlated with σ theta during that block (each dot is data from a separate block from each participant, i.e., the data presented in the middle and bottom rows in a, b); n = 20, directed navigation, purple; n = 16, discovery navigation, green; Spearman *p*=0.01, *r*=0.44). Bottom) Same as above but for μ theta. Here, we only observed a trending correlation with memory performance (Spearman *p*=0.08, *r*=-0.03). **d)** Behavioral performance (color axis) shown as a function of μ and σ of theta difference demonstrating that better memory performance is observed when both μ and σ are low.

We then sought to determine whether memory performance was related to the prevalence of theta oscillations given prior findings showing links between theta and memory^22,23^. To investigate this, we computed the difference in theta prevalence during encoding and retrieval for each recording contact in each participant, and during each block (henceforth referred to as “theta difference”; Methods). This theta difference can be interpreted as a similarity measure, such that values closer to zero indicate comparable theta prevalence during the two phases of the tasks. Within each participant, a closer look at theta difference revealed a curious pattern: the blocks during which theta difference was consistently close to zero across all channels (lowest standard deviation and mean) coincided with the blocks where memory performance was most improved (Fig. 3). This pattern was visible in all participants and during both directed (Fig. 3a) and discovery (Fig. 3b) navigation tasks.

At the group level, memory performance was negatively correlated with the standard deviation of theta difference (Fig. 3c, top; Spearman p = 0.01, *r* = −0.44) and trended towards a negative correlation with the mean (Fig. 3c, bottom; Spearman *p* = 0.08, *r* = −0.30). In other words, as theta prevalence during encoding across all recording channels resembled that during retrieval, memory performance was better. We further quantified this effect by modeling the behavioral performance as a function of both mean and standard deviation of theta difference, as well as an interaction term, using Generalized Estimating Equations (GEEs, Methods). Our results indicated that all three are significant factors in determining behavior (s.d. of theta difference: *p =* 0.001, β = −2.81, Wald χ^2^ = 10.36; mean of theta difference: *p =* 0.023, β = −1.86, Wald χ^2^ = 5.13; interaction term: *p =* 0.021, Wald χ^2^ = 5.35) with a stronger (more negative) modulatory coefficient for the standard deviation. The fact that this effect was observed when considering all of the recording contacts in a participant may indicate a more global theta dynamic effect rather than a local one. For instance, theta prevalence may be similar during encoding and retrieval on one or two contacts (e.g., Figure 3a top, Participant 2, block 3), however, this was not sufficient to predict behavior. The spatial extent of this effect across the brain and the regions recruited during ambulatory spatial memory in humans warrants additional investigation in future studies.

After having established a relationship between theta oscillations and spatial memory, we sought to examine whether theta power exhibited hexagonal modulation with respect to the participants’ head direction during walking. This effect is thought to resemble the grid cell system for navigation in rodents and has been observed in stationary human participants performing stationary 2-D virtual navigation^13,14,24,25^ (Fig. 4a, b; Methods), however, has not yet been explored in freely moving humans navigating a real environment. We found that four contacts (in 3 participants) during discovery navigation and two contacts (in 2 participants) during directed navigation exhibited significant modulation (z > 2) in the theta band but not in the beta or low gamma bands (z < 2; Fig. 4c, d). We found that out of five locations that we hypothesized would exhibit hexadirectional modulation (contacts in the subiculum and entorhinal cortex), three showed significant modulation, whereas only three out of remaining fifteen in other areas showed this modulation (Extended Data Table 2). As expected, normalized theta power was significantly positive when movement was aligned with the preferred orientations and negative otherwise (Fig. 4d, Extended Data Fig. 4; individual participant data), and this was true within each participant (Fig. 4e; Sign-rank test, *p* < 0.05 in each condition).

**Figure 4.**
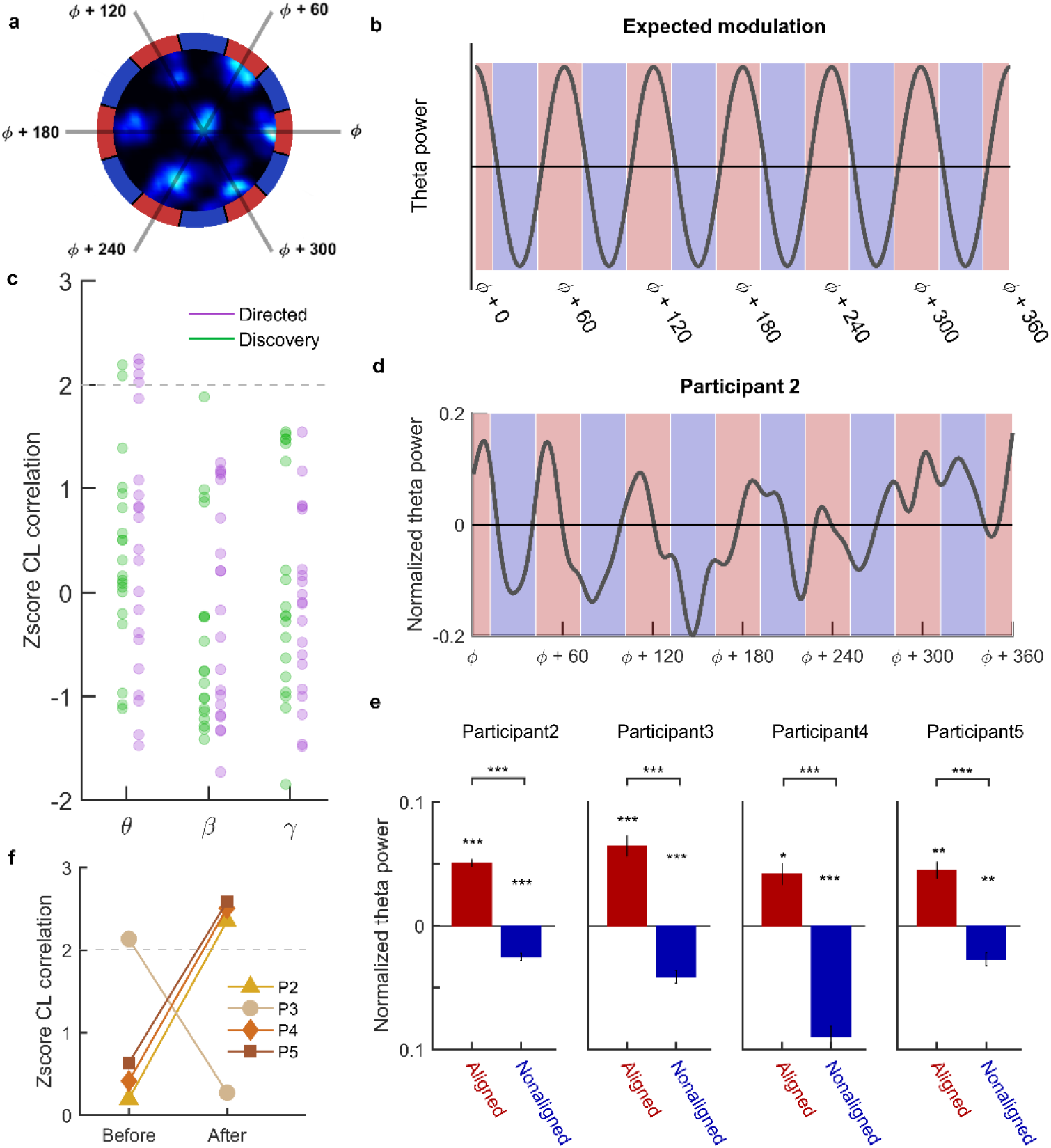
Hexadirectional modulation of theta oscillations as a function of head direction during movement. **a)** Illustration of the phenomenon of a grid cell’s firing rate (lighter shades correspond to higher firing rates) as a function of animal position in an environment. The outer disk demonstrates a six-fold symmetry in neuronal firing with respect to heading direction (red: high; blue: low, here and throughout figure). **b)** Expected modulation of theta power versus head-direction according to (**a**). **c)** Using the circular-linear correlation coefficient methods (Methods), four contacts during directed (purple) and two contacts during discovery navigation exhibited significant hexadirectional modulation of theta (6-12Hz; z > 2), but not beta (13-20Hz) or low gamma (20-60Hz). Dashed gray line indicates z = 2, i.e., significance threshold. **d)** Example data from a participant showing higher theta power during head-directions aligned with the preferred orientation (red) and lower theta power during non-aligned movements (blue) followed a six-fold symmetry. **e)** For the recording contacts with significant hexadirectional modulation, theta power was significantly positive (Wilcoxon sign-rank test) during aligned movements and significantly negative during nonaligned movements for each participant (Extended Data Fig. 4), and significantly different between the two (aligned vs. nonaligned) conditions (Wilcoxon rank-sum test). *p < 0.05, ***p < 0.001. **f)** In three (of the four) participants with significant modulations, the hexadirectional modulation was developed after the behavioral block with high performance (also determined by similarity of theta prevalence during encoding and retrieval, Fig. 3). Dashed gray line indicates z = 2, i.e., significance threshold.

Lastly, we explored whether there was a relationship between memory performance and the emergence of hexadirectional modulation of theta. Grid cell representation in rodents exhibit expansion in scale and reduction in regularity in response to novelty in the environment^26^, which might affect the strength of hexadirectional modulation of theta power with experience. In humans, it has been demonstrated that the hexadirectional representation of theta stabilizes over time in stationary subjects^14^ and the strength of modulation is correlated with behavior^13^ during stationary 2-D virtual navigation. In our data, we first selected the behavioral block during which theta prevalence was similar during encoding and retrieval, which in turn corresponded to the block with better performance (Fig. 3). We then separated the data into blocks before and after this selection point and performed the same analysis to detect hexadirectional modulation of theta power. We found that in three (of the four) participants with significant modulation (found in previously presented analyses during the entire experimental session), the modulation emerged during the second phase of the experiment after learning had occurred (Fig. 4f). We observed the opposite trend in a single participant, however the performance in this participant was overall much lower than the other participants. Altogether, this preliminary finding suggests that spatial learning could influence the strength of the observed hexadirectional theta code for space.

To our knowledge, this is the first study to probe intracranial theta oscillations in the human brain during untethered freely moving navigation with spatial memory demands. We present here a new platform that enables the investigation of neural correlates underpinning naturalistic ambulatory behaviors in humans. Our results reveal that theta oscillations are associated with spatial memory and may serve as a medium to reinstate the memory of previously learned locations. Furthermore, the finding of hexadirectional modulation of theta according to head direction during physical motion, and its relationship with memory, extends previous rodent navigational findings to substrates of human memory in real environments. Taken together, the critical role of theta oscillations in both spatial memory and navigation in humans may further link the neural mechanisms underlying these seemingly disparate functions.

## Acknowledgments

We thank FabLab (Golden Beach, FL), Interactive Lab (Moscow, Russia), and Merit Vick (NeuroPace, Inc.) for technical assistance. We thank Thomas Wolbers for useful advice and discussions. This work was supported by UCLA startup funds and the National Institute of Neurological Disorders and Stroke (NS103802; NS058280). We also thank the participants for taking part in our study.

## Materials and Methods

### Data Acquisition

Intracranial EEG data acquisition and motion tracking were done using previously described methods (M. Aghajan *et al.*, 2017). In brief, the FDA approved NeuroPace RNS System was used to record iEEG activity from depth electrode contacts (1.27 mm diameter; each with 4 platinum-iridium electrode contacts; surface area = 7.9 mm^2^, length = 1.5 mm; electrode spacing = 3.5 or 10 mm; Extended Data Table 2) chronically implanted in patients with pharmaco-resistant epilepsy. The RNS System records iEEG from four bipolar channels (sampling frequency = 250 Hz; analog bandpass filter, cutoff frequencies: [4 − 90] Hz, attenuation: 3 db). The blind participant is implanted with the newer RNS-320 System, upgraded since our previous study, for which the cutoff frequencies were clinical default settings [5 – 70] Hz. A custom-made Electromagnet was externally placed on the head over the position of the underlying implanted RNS System and was used to trigger data storage and simultaneously generated a visible light signal detected by the motion capture system that was then used to then synchronize iEEG and behavioral movement data. Due to variable, and uncontrollable, delays in the magnet signal (< 500ms)^27^, the synchronization between behavior and neural data only allowed for analyses that required lower temporal precisions (e.g., phase of the experiments such as encoding/retrieval rather than trial-based analyses).

### Task Implementation

To achieve room-scale virtual spatial navigation, we combined motion capture and VR technologies. Motion capture was done using the Optitrack system (Natural Point, USA) with 12 ceiling mounted cameras; 10 cameras were configured to track a rigid body (a unique configuration of infrared reflective markers located on the participant’s head), and 2 additional cameras were allocated to record video for synchronization purposes. The paradigms were implemented in the Unity game engine and the applications were built for a Samsung Galaxy S7 device that was used with a Samsung GearVR goggles (both of which were products of SMI; sampling rate = 60 Hz).

The VR applications were also configured to communicate with the motion capture system via WiFi through custom Unity scripts (Interactive Lab, Russia), thus allowing for real-time room scale virtual navigation, *i.e.*, the physical movement of the participant in the real room, tracked with the motion capture cameras, was mapped to their position in the virtual room, and as they moved in real-time and thus the virtual scene was updated in a one-to-one manner. A wireless joystick was paired with the headset phone (that ran the VR application), and was used to track button presses during the retrieval phase of the experiments (see Behavioral tasks section below).

### Behavioral Tasks

The behavioral tasks consisted of three distinct phases: 1) encoding, 2) distraction, and 3) retrieval. During the encoding phase of each block, participants were instructed to learn the locations and the order of multiple yellow translucent virtual cylinders (halos; height = 120 cm, diameter = 40 cm, for a detailed description, see below). This was followed by a distraction phase, during which participants were asked to count backwards from 100 by three for a duration of 30 seconds. This non-mnemonic task was chosen to increase the long-term memory demands of the spatial navigation task and thus its hippocampal dependency. During the retrieval phase, participants were instructed to navigate to the locations of the previously learned halos, in the order that they were originally visited, and to press a button on a handheld wireless joystick at the moment of arrival.

Each participant performed the same task with fixed locations over several alternating blocks (4-6) of encoding, distraction, and retrieval. Instructions were presented through the VR headset at the beginning of each experimental phase. Additionally, before data collection, there was a practice session during which participants performed the same task but in a different virtual environment to ensure they became familiar with the task structure and could comfortably navigate but not exposed to the to-be-learned environment (Extended Data Fig. 1).

#### Ambulatory virtual spatial navigation (sighted participants)

After a practice session, participants completed both a directed and discovery version of the spatial navigation task, with the order of task versions counterbalanced across participants. Further details of each task are provided below:

1. Directed navigation task: During encoding, participants were instructed that they would see several (10 locations) visible halos that would appear one at a time in a different location within the room (halo locations were predetermined to maximize the distance between two consecutive halos). Upon seeing a halo, they were asked to walk towards it, stand in the center until it disappeared (after 4 seconds), and remember the location and the order in which the halos were presented. After the distraction phase, participants were provided with a joystick and asked to physically navigate to the center of the halos that they previously visited during the encoding phase, in the order they learned them, and press a joystick button upon arrival. Retrieval phase had a fixed duration of 60 seconds (Supplementary Video 1).
2. Discovery navigation task: Task structure closely resembled the directed navigation task with the exception of how halos were presented. Here, participants were instructed to freely explore the VR room and find multiple halos (6 locations) that would appear one at a time and only when participants were in close proximity to them (1 meter away from the halo center and remained visible until exiting. Here, too, they were asked to remember the presentation order of the halos in addition to the locations. The retrieval phase here was the same as in the directed navigation task but only lasted 45 seconds, since there were fewer locations (6) to be remembered compared to the directed navigation task (10) (Supplementary Video 2).

#### Ambulatory virtual spatial navigation (blind participant)

The blind participant performed an auditory version of the tasks described above. Here, the location of the halos in the directed navigation task were marked with a continuous auditory cue (beeping sounds) that increased (decreased) in frequency (i.e., pitch) as the participant approached (moved away) from the halo locations. For the encoding phase in the discovery navigation task, the sound only initiated upon arrival 1 meter from the halo centers. During retrieval phase for both tasks, the participant explored the room using a hoover cane in one arm and a hand gesture with the other to signify arrival upon halo locations.

### Behavioral performance metrics

Memory performance on the directed and discovery navigation tasks was calculated using two measures during retrieval: (1) accuracy in position, i.e., how far away the recalled halo locations were to the original halo locations presented during encoding; (2) accuracy in order, i.e., how closely the order of the recalled halo locations resembled the true order presented during encoding. To measure the accuracy in position, we constructed a Gaussian probability distribution centered around the center of each halo, and with a variance equal to the halo diameter. The maximum value of the curve was then mapped to 1, and the location of each recalled halo was scored based on this curve (a final normalization bound this metric in the range [0, 1]). To quantify the performance in terms of order, two criteria were taken into consideration: (1) the visited recalled n^th^ halo correctly corresponded to the encoded n^th^ halo (e.g., 4^th^ halo during encoding was visited 4^th^ during retrieval regardless of the order of the halos before or after during retrieval); (2) n^th^ and n+1^th^ halo were visited in order (e.g., 4^th^ and 5^th^ encoded halos were visited in sequence during retrieval, even if they were the first two button presses). These correspond to the sum over the diagonal and off-diagonal elements of a binary n × n matrix consisting of encoded (rows) and retrieved (columns) halo locations. The overall memory performance was the average value of the position and order scores.

### Electrode Localization

A high-resolution post-operative CT image was obtained and co-registered to a pre-operative whole brain and high-resolution MRI for each participant using previous methods^28^ (Fig. 2a; Extended Data Fig. 2). Segmentation of MTL regions (entorhinal, perirhinal, parahippocampal, hippocampal subfields CA23DG [CA2, 3, dentate gyrus], CA1, and subiculum) was done using the automated ASHS software^29^.

### Processing iEEG data recorded via the RNS System

The processing of the iEEG data (sampling rate = 250Hz) was similar to recently published methods (M. Aghajan et al., 2017). However, these methods are briefly summarized below:

i. *Detection and Elimination of epileptic activity.* Putative epileptic activity was detected using thresholding algorithms^8,30^ and functions found in the MATLAB signal processing toolbox. Thresholding involved two criteria and one of them needed to be met in order for the detection to occur: 1) the envelope of the unfiltered signal was 5 s.d. above the baseline; and 2) the envelope of the filtered signal (25-80Hz band-pass filter after signal rectification) was 6 s.d. above the baseline. This was then followed by visual inspection of the data. Epileptic epochs in the iEEG (median, [25^th^, 75^th^] = 2.92%, [2.33%, 3.85%] and their corresponding behavioral epochs, were discarded from further analysis.
ii. *Calculation of oscillatory power and prevalence.* We used the BOSC toolbox^21,31^ to perform time-frequency analysis as well as detect episodes with significant oscillations at any frequency (sixth order wavelets were used and bouts were required to occur for at least three cycles and above 95% chance level). Prevalence of an oscillation (theta in our case) was defined as the percentage of time (compared to the total time duration) with significant oscillations in the 6-12Hz frequency range.
iii. *Extraction of theta activity from iEEG data.* We applied an acausal Butterworth filter of the fourth order to bandpass the raw signal in the 6-12Hz frequency band (Fig. 2b).

### Theta prevalence and behavior

The goal of this analysis was to examine whether there was a relationship between specific features of the theta oscillation and spatial memory performance.

i. *Theta similarity between encoding and retrieval.* For each participant, theta prevalence (described in the previous section) was computed for each recording contact (n_contacts_ = 4), for all behavioral blocks (n_blocks_ = 4-6, depending on the participant and task), and during each phase of encoding (Θ_encoding_) and retrieval (Θ_retrieval_) separately. Thus, an element-wise subtraction in theta prevalence during encoding and retrieval resulted in a matrix of size (n_contacts_ × n_blocks_), which was then normalized by the sum of the theta prevalence (Fig. 3a, b; top rows). This quantity is referred to as “theta difference” and is defined as: (Θ_encoding_ – Θ_retrieval_)./ (Θ_encoding_ + Θ_retrieval_). We then computed the mean and standard deviation of theta difference across the recording channels (Fig. 3a, b; middle rows). These values (mean and s.d. of theta difference) was subsequently correlated with the participants’ behavioral performance during each block.
ii. *Generalized Estimating Equations (GEEs).* To assess the effect of theta prevalence (namely the mean and standard deviation of theta difference as a measure of similarity) on behavioral performance during each block, we used GEEs. This approach allows for handling potential clustering of the data points due to the within-subject repeated measure design. Here behavioral performance was modelled as a function of the mean and s.d. (of theta) covariates as predictors (also including the interaction term, and an intercept), participant numbers as the subject variable, with linear scale response, identity link function, and exchangeable correlation type. The reported statistics (found in the main text) include the p values, Wald χ^2^ statistic, as well as estimated β coefficients. This analysis was done using SPSS software (IBM Corporation, USA).

### Hexadirectional modulation of theta power

We employed methods that were previously applied to fMRI data from immobile participants^15^ and were recently adapted to human iEEG data also in immobile participants who completed a view-based 2-D VR spatial navigation task^13,14^. First, we computed the allocentric head-direction of the participant (defined as the orientation of the rigid body tracked with the motion capture system) with respect to the physical room. Head-direction data (sampled at 120Hz) was interpolated to match the iEEG data in time. We then computed the oscillatory power in 3 distinct frequency bands of theta (6-12Hz), beta (12-30Hz), and low gamma (40-60Hz) (high gamma range of 60-100Hz could not be achieved due to the 90Hz upper limit of the bandpass filter on the RNS System). For these analyses, only data during movement (speeds above 10cm/s) were included.

a. *Circular-Linear correlation method*. In this method, a correlation coefficient (ρ, and its corresponding p value) was computed between the head-direction of each participant (circular variable) and the oscillatory power (linear variable) at any given time. This was done using the circular statistic toolbox^32^. Furthermore, we generated surrogate data by circularly shifting the iEEG data with respect to the head-direction (n = 500) to calculate a z-score value for the correlation coefficients for a given electrode in each participant.
b. *Generalized Linear Model (GLM) Method.* We divided the iEEG data (during both encoding and retrieval phases) into a training (two-thirds of the data) and testing (one-third of the data) set. The training data was used to find the preferred grid orientation (θ_0_), which was later used to calculate the hexadirectional modulation index on the test set. The preferred grid orientation was found by applying a GLM fit as follows:

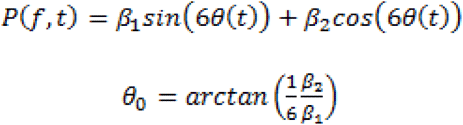

Where θ(*t*) and *P*(*f,t*) are the head-direction and oscillatory power at a given frequency range at time t. Once θ_0_ was found, the head-direction was adjusted (θ′(*t*) − θ(*t*) − θ_0_) and a β score was computed by applying another GLM fit on the test data:

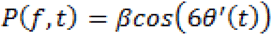 This method was used to determine the grid preferred orientation that was later used to compute the modulation of theta power during aligned and nonaligned movement directions (part c). However, the results from this part of the analysis, i.e., computing betas and their associated statistic were not reported here.
c. *Binning methods (theta power along movement directions).* Theta power (total power within the 6-12Hz) was first z-scored. The true head-direction of the participant (the output of the motion capture system) was reset to match the preferred grid orientation (found using the GLM method). The new directions were then binned into 1 degree angles and z-scored theta power was averaged in each direction bin, and smoothed (using a Gaussian kernel of 4 degrees). The results from this analysis are shown in Fig. 4d and Extended Data Fig. 4. In order to obtain theta power during aligned versus nonaligned movements (Fig. 4e), we used a similar approach. Here, movement directions were considered aligned with the preferred grid orientation if they were within 30 degrees angles around [0, 60, 120, 180, 240] degree directions, and nonaligned if otherwise.

---

**Extended Data Figure 1.**
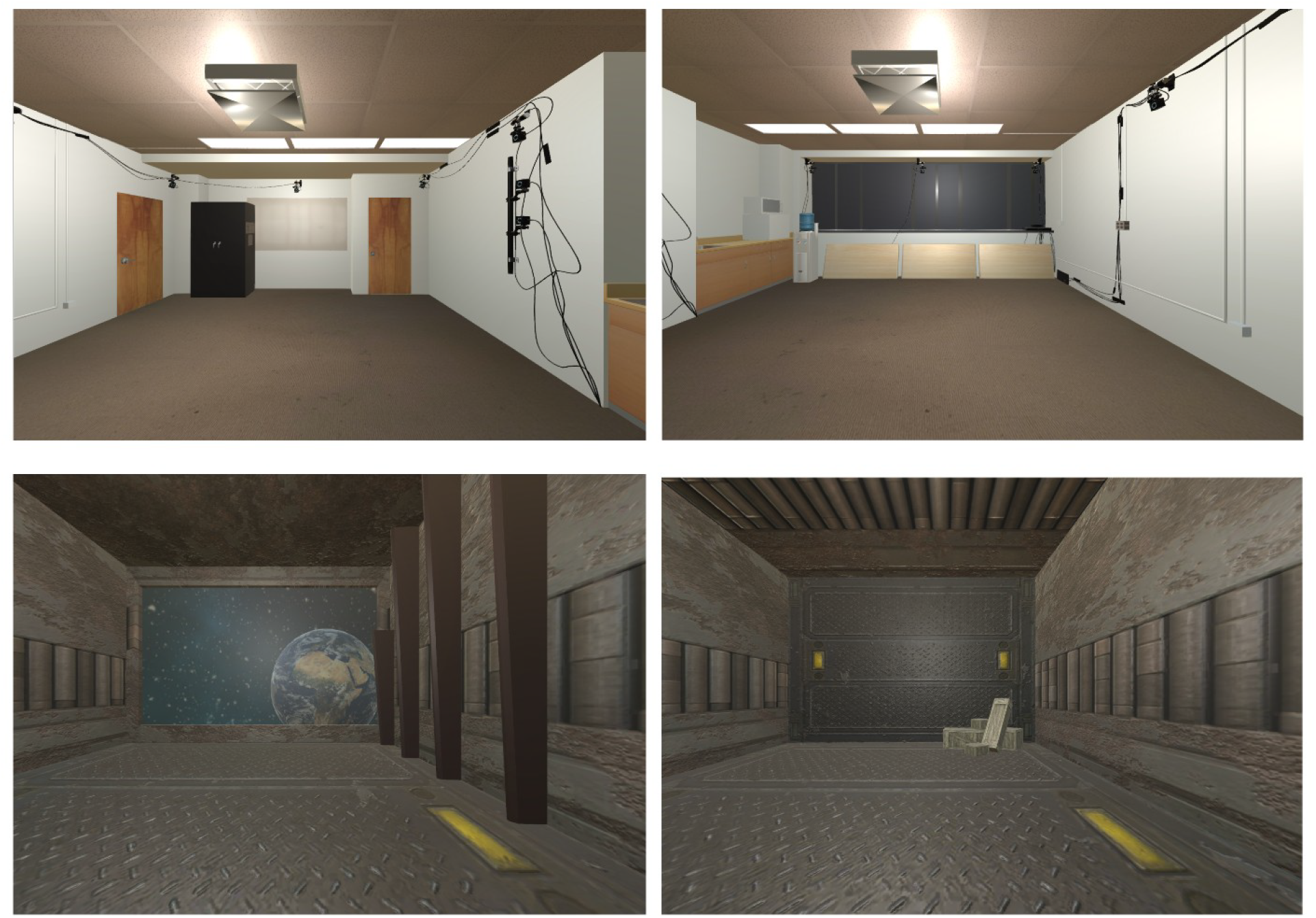
Virtual rooms. Top row) North and south views of the virtual environment in which the participants performed the directed and discovery spatial navigation tasks. Bottom row) North and south view of a second virtual environment that was used for the pre-testing practice navigation session prior to completing the experimental tasks.

**Extended Data Figure 2.**
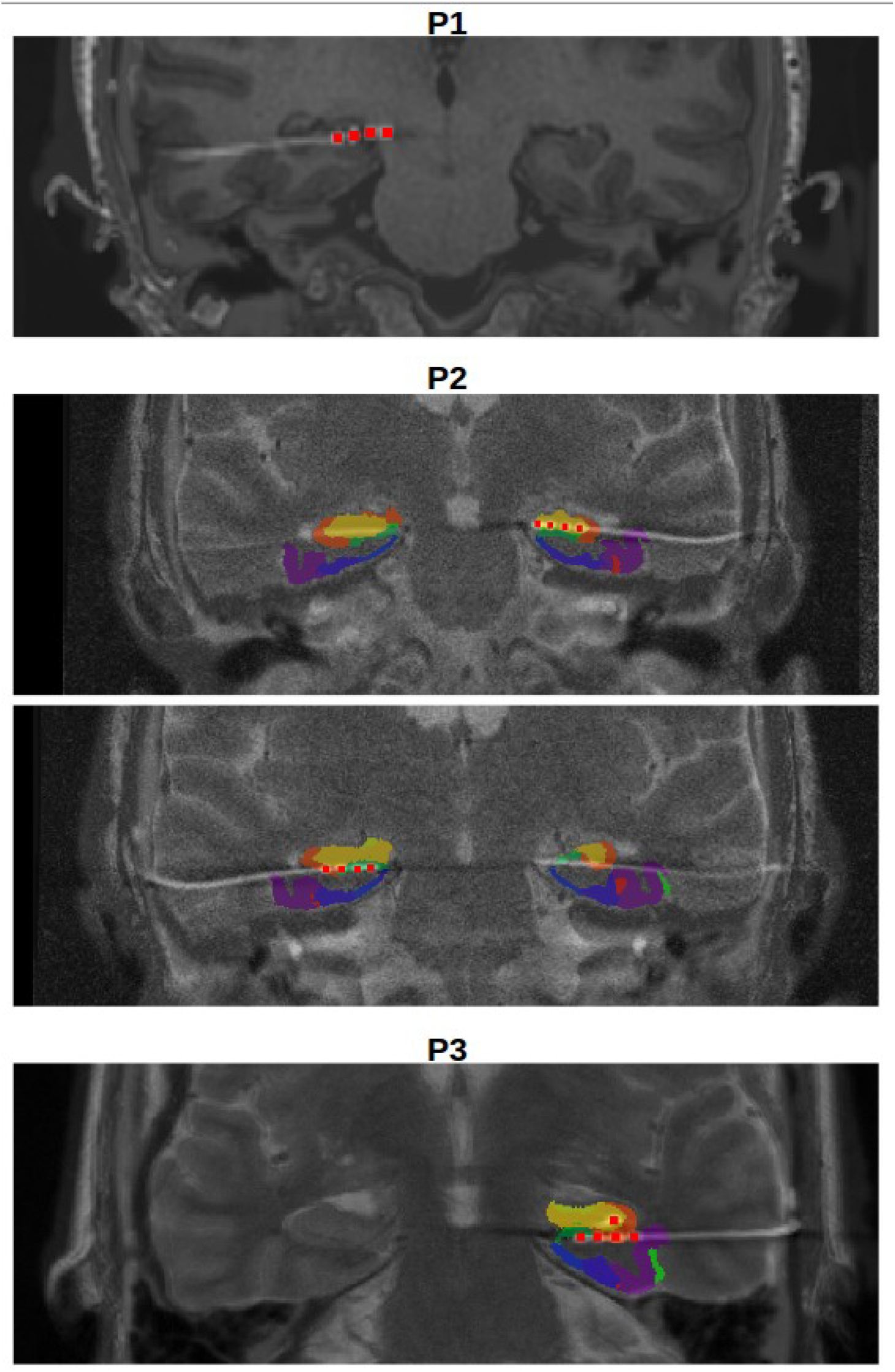
Example electrode localizations from all participants. Electrode locations are marked in red squares. Different colored areas are the results of an automated hippocampal segmentation using ASHS software. A high-resolution T2 MRI was not available for participant 1 (P1) and thus electrode locations are shown without overlaying MTL subregional segmentations. Electrode locations for P4 are shown in main Figure 2, and for P5 were previously published in M. Aghajan *et al.* (blind participant)^8^. For a complete list of electrode locations see Extended Data Table 2.

**Extended Data Figure 3.**
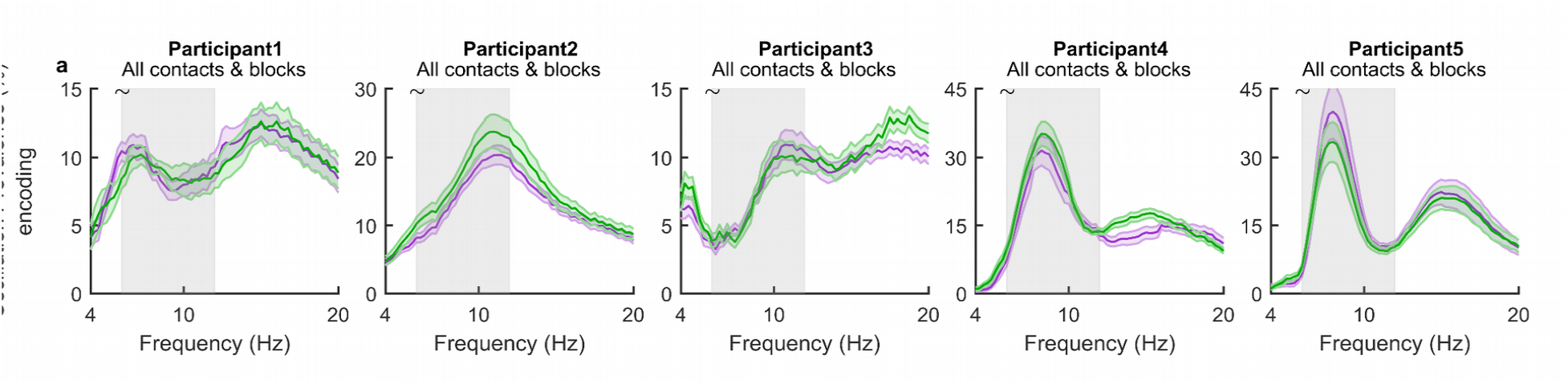
Theta prevalence was similar during the retrieval phase of directed and discovery navigation tasks. **a)** Theta prevalence was similar (*p*>0.05; clustered-based permutation test) between directed (purple) and discovery (green) navigation in all participants (when considering all channels and behavioral blocks; P1: n1=20, n2=16; P2: n1=24, n2=20; P3: n1=20, n2=8—data was only collected during blocks 3-4; P4: n1=20, n2=16; P5: n1=20, n=16). Shown are the mean (dark color lines) ±s.e.m (shaded color regions) in each participant. Gray area indicates the bounds of the theta frequency band chosen (6-12Hz).

**Extended Data Figure 4.**
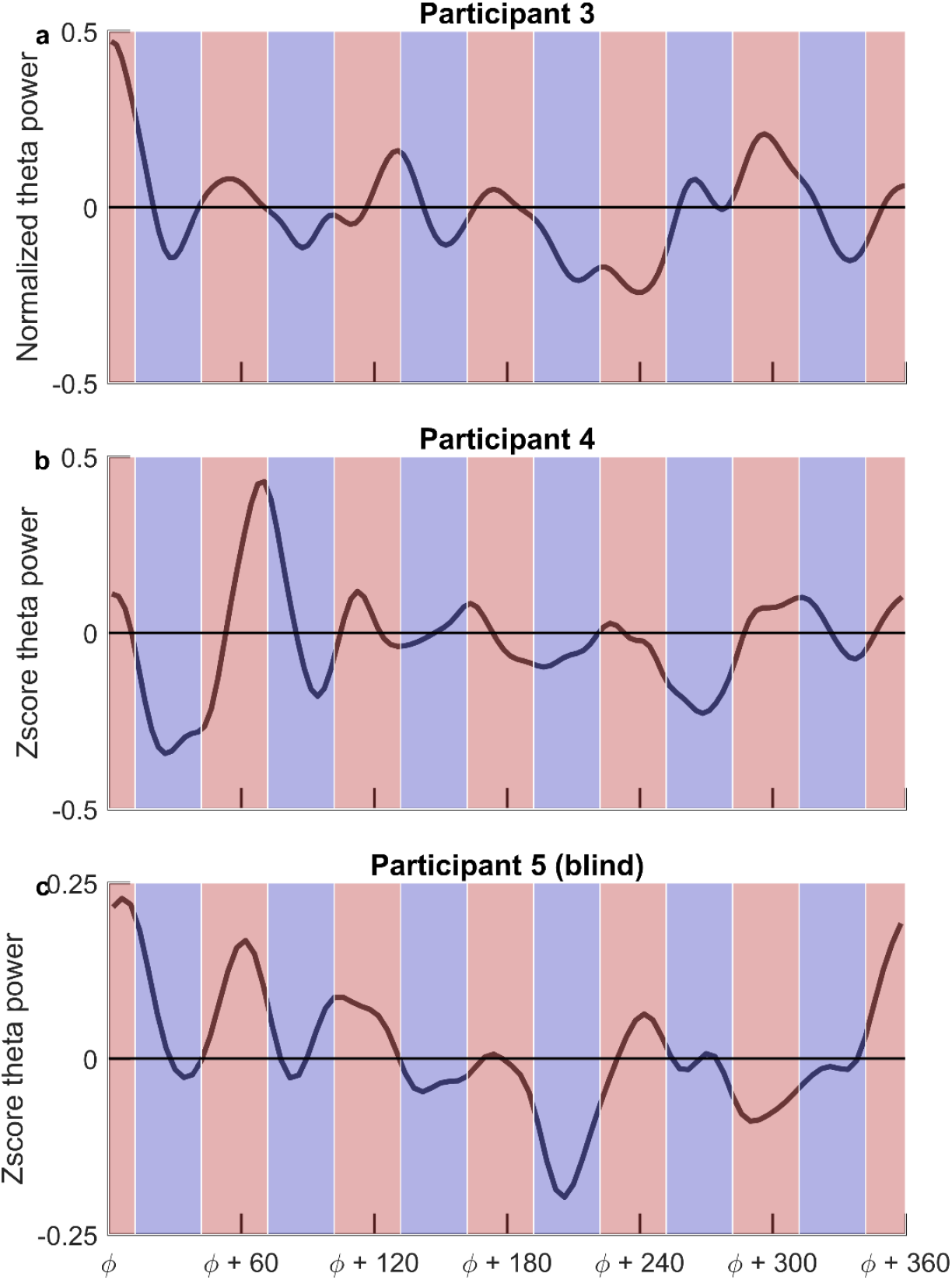
Additional participant examples of hexadirectional modulation of theta power as a function of head-direction. Theta power was z-scored and binned according to the participants’ movement direction. Red and blue areas correspond to directions that were aligned and non-aligned with the preferred grid orientation respectively.

**Extended Data Table 1.** Participant demographics. Demographics of the study participants including age, gender and handedness. * Participant P5 is congenitally blind.

| Participant | Age | Gender | Handedness |
| --- | --- | --- | --- |
| P1 | 21 | M | R |
| P2 | 40 | M | R |
| P3 | 51 | F | R |
| P4 | 48 | F | R |
| P5* | 65 | M | R |

**Extended Data Table 2.** Electrode Localizations. Electrode labels were determined based on clinical criteria and include R(L)EC, corresponding to the right (left) entorhinal cortex, R(L)HIP corresponding to the right (left) hippocampus, and one participant who had ROFC (right orbitofrontal cortex) contacts. The precise location of clinical electrode labels determined after placement using high-resolution MRI/CT are shown in columns 4 and 5^28^ (Fig. 2a, Methods). Lower digits in the electrode contact labels indicate more distal contacts. Recordings were bipolar using adjacent electrodes 1-2 and 3-4 (e.g., REC1-REC2 and REC3-REC4 are two example iEEG channels). Shaded cells correspond to the contacts on which significant hexadirectional modulation of theta power with respect to head-direction was observed during discovery (green) and directed (purple) navigation tasks (Fig. 4).

| Subject | Contact Labels | Electrode Spacing | Localization 1 | Localization 2 |
| --- | --- | --- | --- | --- |
| P1 | RHIP1 – RHIP2 | 3.5 mm | -- | Subiculum |
|  | RHIP3 – RHIP4 |  | -- | CA1 |
|  | ROFC1 – ROFC2 | 10 mm | Orbitofrontal Cortex | Orbitofrontal Cortex |
|  | ROFC3 – ROFC4 |  | Orbitofrontal Cortex | Orbitofrontal Cortex |
| P2 | LAH1 – LAH2 | 3.5 mm | CA23DG | CA23DG |
|  | LAH3 – LAH4 |  | CA23DG | CA23DG |
|  | RAH1 – RAH2 | 3.5 mm | Subiculum | Subiculum |
|  | RAH3 – RAH4 |  | CA23DG | CA1 |
| P3 | LHIP1 – LHIP2 | 10 mm | Subiculum | Subiculum/CA1 |
|  | LHIP3 – LHIP4 |  | CA1 | Perirhinal White |
|  | LEC1 – LEC2 | 3.5 mm | Amygdala | CA1 |
|  | LEC3 – LEC4 |  | CA23DG | CA23DG/CA1 |
| P4 | LEC1 – LEC2 | 10 mm | Hippocampus | Perirhinal Cortex |
|  | LEC3 – LEC4 |  | Inferior Temporal | Inferior Temporal |
|  | RAH1 – RAH2 | 3.5 mm | Subiculum | Subiculum/CA1 |
|  | RAH3 – RAH4 |  | CA1 | Perirhinal White |
| P5 | LEC1 – LEC2 | 10 mm | Entorhinal Cortex | Perirhinal Cortex |
|  | LEC3 – LEC4 |  | Inferior Temporal | Inferior Temporal |
|  | LHIP1 – LHIP2 | 3.5 mm | CA1 | CA1 |
|  | LHIP3 – LHIP4 |  | CA1 | CA1 |

